# The Neural Basis of Predictive Pursuit

**DOI:** 10.1101/694604

**Authors:** Seng Bum Michael Yoo, Jiaxin Cindy Tu, Steven T. Piantadosi, Benjamin Yost Hayden

## Abstract

It remains unclear how and to what extent non-human animals make demanding on-the-fly predictions during pursuit. We studied this problem in a novel laboratory pursuit task that incentivizes prediction of future prey positions. We trained three macaques to perform joystick-controlled pursuit of prey that followed intelligent escape algorithms. Subjects reliably aimed towards the prey’s likely future positions, indicating that they generate internal predictions and use those predictions to guide behavior. We then developed a generative model that explains real-time pursuit trajectories and showed that our subjects use prey position, velocity, and acceleration to make predictions. We identified neurons in the dorsal anterior cingulate cortex (dACC) whose responses track these three variables. These neurons multiplexed prediction-related variables with a distinct and explicit representation of the prey’s future position. Our results provide a clear demonstration that the brain can explicitly represent future predictions and highlight the critical role of anterior cingulate cortex for future-oriented cognition.

**One-sentence summary:** In a dynamic pursuit environment, monkeys actively predict future prey positions and dACC neurons encode these future positions.

## INTRODUCTION

Many foragers pursue fleeing prey. The ability to effectively pursue prey is thus a critical element in our behavioral repertoires ^1, 2^. To pursue effectively, a forager needs to perform a series of computations: it must maintain a representation of its current position relative to that of the prey, then compute a best path to capture the prey, then execute that path. Because the ability to perform such computations can determine foraging success, pursuit has likely been an important driver of our cognition and its underlying brain systems ^3–6^.

One way to improve pursuit effectiveness is to predict the future position of the prey and head towards the predicted position ^7^. Estimating future positions can be done using the prey’s basic Newtonian variables (most importantly, its current position, velocity, and acceleration) and can be improved using additional (potentially even recursive) variables, such as predictions about the likely evasive strategy of the prey in response to the predator’s own future path. By using such information, the forager may be able to formulate a representation of the predicted future position of the prey. The ability of non-human animals to actively predict positions of prey during pursit is poorly understood. Nonetheless, predictive pursuit is an important part of the repertoire of many species.

Prediction is important for many cognitive and behavioral processes, not just foraging. These include motor control, economic decision-making, and abstract long-term planning ^8–14^. There is some evidence that foraging animals can predict the long-term future - that is, they may be able to travel mentally in time and see themselves in the future ^15, 16^. However, observations about animal prediction tend to be limited to a small number of highly adapted species in unique contexts. And, while future planning of movements is relatively well-studied, the ability to predict future positions of prey during dynamic behavior with rapidly changing goals – which feed into but are distinct from motor plans – is not. In the context of pursuit, a critical question is whether future-predicting foragers maintain a specific representation of potential future prey positions and whether those representations (assuming they exist) make use of specialized processes.

Although the neural bases of predictive pursuit remain unclear, we can draw some inferences about its likely neuroanatomy. In particular, the dorsal anterior cingulate cortex (dACC) has been implicated in prediction, prospection, and related processes ^17–20^. For example, neuroimaging studies indicate that human dACC is a key region for economic prediction ^21^, for prospective reasoning ^11^ and for more open-ended prospective processes ^21, 22^. The dACC is well-positioned for this role: it receives broad inputs from limbic and cognitive systems, integrates these, and generates high-level control signals that regulate behavior in an abstract and high-level way ^19,22–24^.

Here, we examined the future predicting abilities of rhesus macaques using a novel *virtual pursuit task*. Subjects used a joystick to move an avatar in an open two-dimensional field displayed on a computer screen. Subjects, controlling the avatar, pursued a fleeing prey item that used an artificially intelligent (AI) algorithm to avoid predation. By examining the properties of a generative model fit to our data, we found that our subjects moved towards extrapolated future positions of prey rather than just pointing towards the preys’ present positions. Our subjects’ made their predictions based on three Newtonian variables associated with the current state, but not other factors that could further improve predictions (such as the effect of the subject’s movements on the future position of the prey). We also found that neurons in dACC were selective for those three Newtonian variables (and not others), indicating that responses in this region provide sufficient information to generate the types of predictions our subjects made.

Finally, we found that dACC neurons used a spatial code to explicitly represent the predicted future position of the prey, and that this future representation is multiplexed with the representation of current Newtonian variables.

## RESULTS

### Behavioral results

Three macaques (*Macaca mulatta,* subjects K, H, and C) used a joystick to control the position of an avatar (a yellow or purple circle) moving continuously and smoothly in a rectangular field on a computer screen (Figure 1 and **Methods**). On each trial, subjects had up to 20 seconds to capture a prey item (a fleeing colored square) to obtain a juice reward. Prey avoided the avatar with a deterministic strategy that combined repulsion from the subject’s current position with repulsion from the walls of the field. The prey item was drawn randomly from a set of five, identified by color, that differed in maximum velocity and associated reward size.

**Figure 1.**
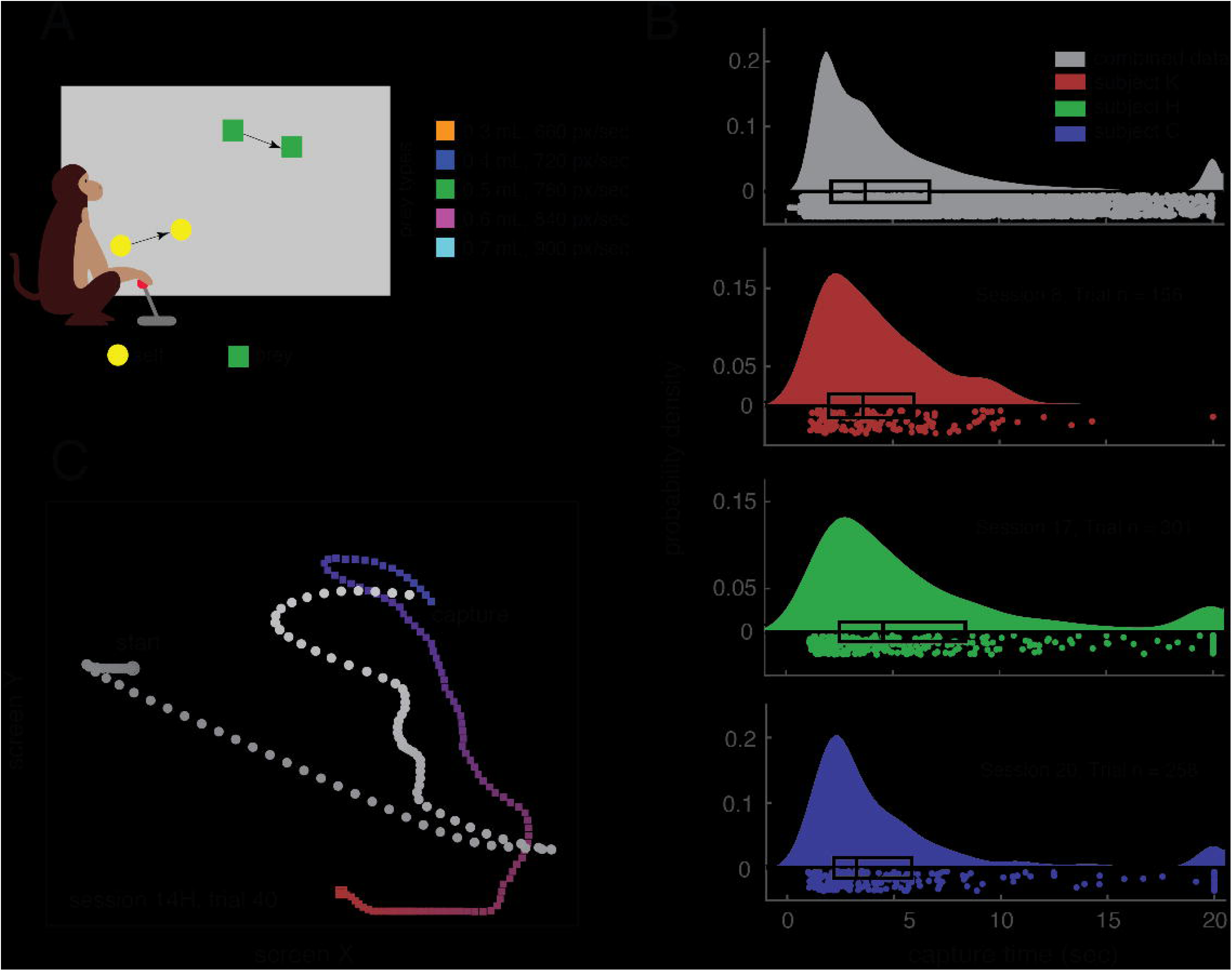
Experimental paradigm and behavioral results. (**A**) Cartoon of *virtual pursuit task*. Subject uses a joystick to control an avatar (circle) and pursue prey (square) on a computer screen. (**B**) Raincloud plot showing each subject’s capture times in an example session (limit was 20 seconds). The box plot indicates 2nd and 3rd quartile of the data; midline indicates the median of the data (K: 3.36 sec, H: 3.73 sec, C: 3.93 sec). The dots under the probability density functions indicate individual data points. (**C**) Avatar and prey trajectories on example trials. Grey: path of avatar; red/blue: path of prey. Color gradient indicates the time progression through the trial.

All subjects showed stable behavior within twelve 2-hour training sessions that followed a longer training period on joystick use (Figure S1 and Figure S2). All data presented here were collected after the training sessions (number of trials, K: 3229; H: 3890; C: 2512). Subjects successfully captured the prey in over 95% of trials and, on successful trials, did so in an average of 5.04 seconds (K: 4.26 sec, H: 5.32 sec, C: 5.54 sec) and median of 3.62 seconds (K: 3.36sec, H: 3.73 sec, C: 3.93 sec). Subjects’ performance varied lawfully with prey type, indicating sensitivity to manipulation of reward and/or difficulty (Figure S1).

**Figure 2.**
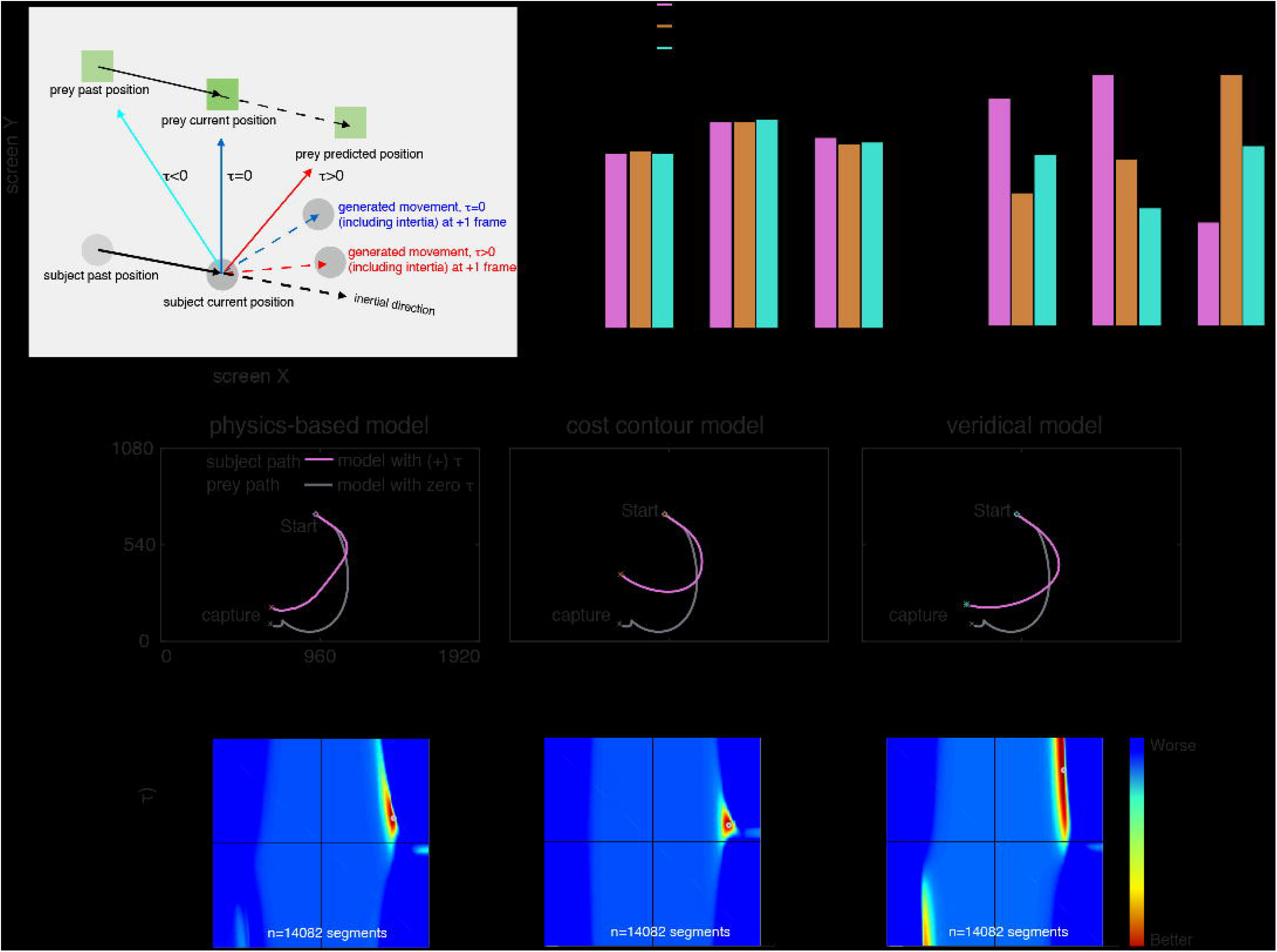
Model description and fitting results. (**A**) Cartoon of model for generating future position based on prediction. Solid black arrow indicates movement from previous time frame to the current one. Subjects are assumed to aim at a point that leads (red solid arrow) or lags (cyan solid arrow) the prey. The resulting movement (red/blue dashed arrow) vectors are constrained to a maximum speed and inertia (black dashed arrow). (**B**) Fitting results: Akaike Information Criterion (AIC, left) across all the trajectories and percentage of trials best explained by each model (right). For calculating the AIC, we summed the log-likelihood across the whole data set from each subject individually and used the quantity (2 x number of the segments) as the number of free parameters. This quantity was: subject K, 28,164; subject H, 35,308; subject C, 20,720 parameters. Predictive models provide better fits than zero prediction ones. **(C)** Example trajectories and corresponding fit trajectories generated by predictive and non-predictive models. **(D)** Heatmap plots of model performance explaining subject’s pursuit segment across parameter space from a single subject (Subject K) for physics based model (left), cost model (center), and veridical (right). The small gray circle at the peak indicates the best parameter combination explaining that subject’s behavior, that is, the one that generates the smallest distance between the actual segment and model-predicted segment.

### Behavioral evidence of future state prediction

For analysis purposes, we split all data into one-second segments (Figure S3). Within each segment, we calculated the error (sum of squares) between the model (see below) and the behavior at each frame (i.e., each 16.67 ms). For each segment, we computed the minimum point on a 201×201 matrix of intensities for each parameter pair (force by time, Figure 2, see below). We then averaged over all segments and all trials, separately for the three subjects.

**Figure 3.**
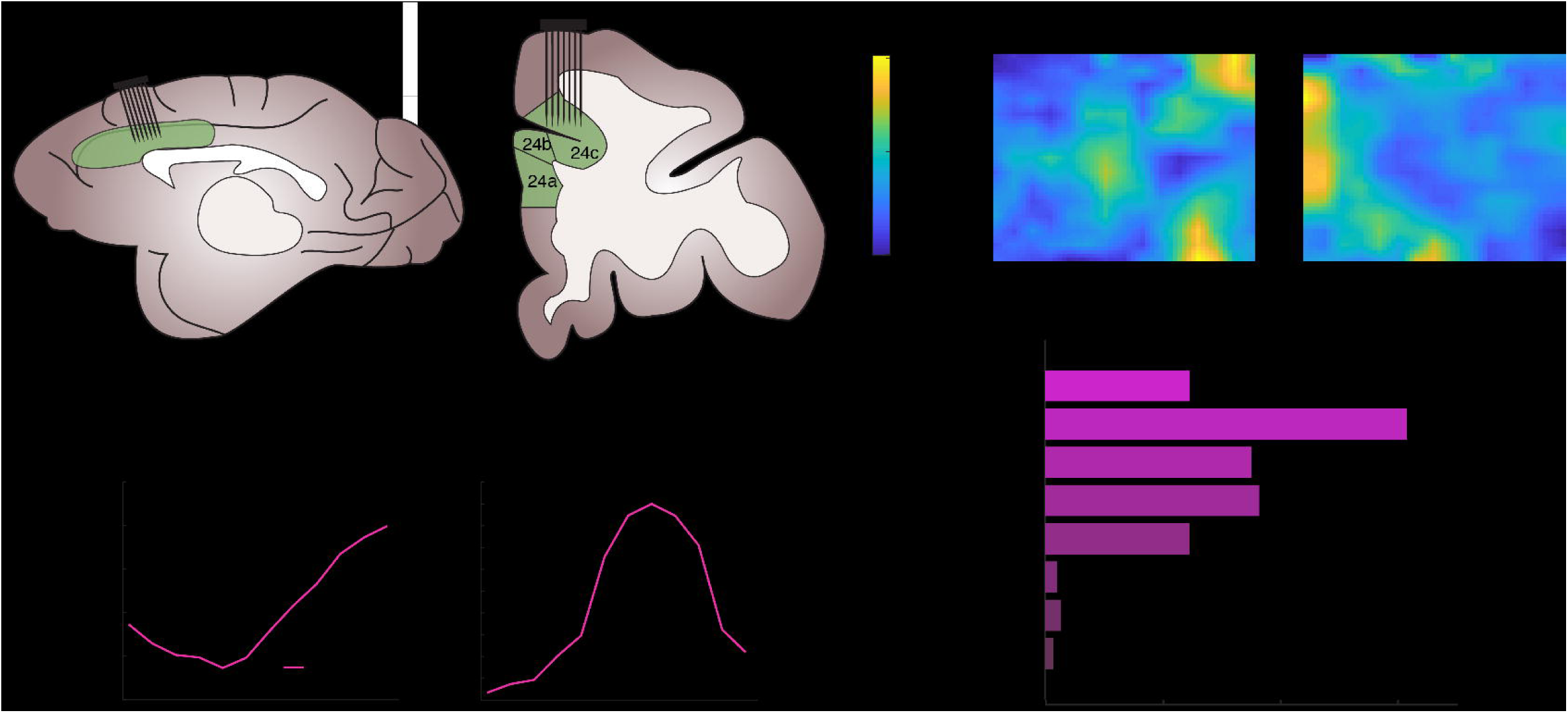
Basic neural results. (**A**) Cartoon showing location of recorded brain areas in dACC; sagittal and coronal views. (**B**) Filters (tuning surfaces) of two example neurons showing selectivity for current position of the prey. (**C**) Example neuron showing tuning for speed (black line) and the corresponding model fit (magenta line). (**D**) Example neuron showing tuning for prey direction (black line) and the corresponding model fit (blue line). (**E**) Preponderance of tuning for the Newtonian physics variables tested. Tuning for future position is counted only if the neuron is selectively tuned for future position above and beyond current position.

We developed a generative model of behavior (see **Methods**). We used the variable τ (tau) to refer to the prediction parameter for each subject. The variable τ comes from the model and refers to a fit scalar variable, which is multiplied by future position (see the equations in **Methods** section “**Behavioral Model**”). In practice, it can be interpreted as the distance into the future that the subject prospects to guide his behavior (Figure 2A). The variable τ can have positive, negative, or zero values. A positive value for τ indicates that the subject points towards the expected future position of the prey - that is, the strategy reflects prediction. A zero τ indicates that the subject points the joystick directly at the current position of the prey. A negative value for τ indicates that the subject points the joystick towards where the prey was in the recent past. Note that all of these strategies (within limits) are capable of eventually catching all prey, since the subject’s avatar is, by design, faster than the prey. The scalar parameter κ (kappa) reflects the amount of force applied toward the direction of the predicted position. Thus, a negative value indicates that force is exerted away from (180 degrees opposite) the prey’s position, whereas a positive value indicates that force is exerted towards it.

We also added an inertia term to the model. Specifically, we computed an inertially biased path for each 16.67 ms frame. The biased path is a vector sum of the computed best predicted direction and the previous direction (P_subject_(t) – P_subject_(t-1)). In our implementation, these two terms have equal weighting. Note that in practice, their relative weighting may nonetheless vary because the force term (κ, which is fit in the model), affects the weight of the new direction relative to the past direction. This approach for implementing inertia is designed to align intuitively with how inertia works (see **Methods**, Figure S2, and S4).

**Figure 4.**
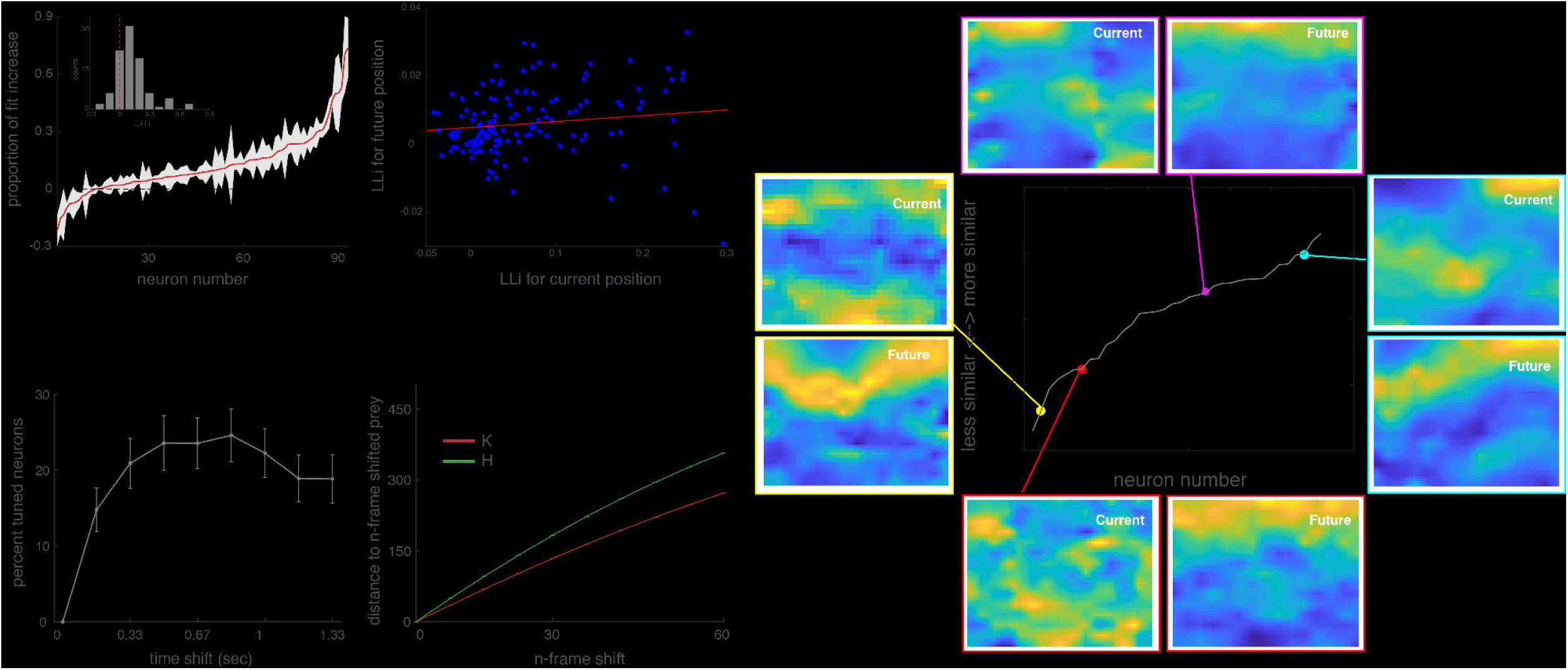
Properties of future position selectivity. **(A)** Proportion of variance explained by including future position in each neuron (only neurons that are selective for current position are shown). Neurons are sorted according to amount of additional variance explained by future position. **(B)** Log likelihood increase (LLi, a measure of explanatory power) for current and future position are correlated on a cell-by-cell basis. Red solid line indicates the linear regression line. **(C)** Example filters from neurons that are significantly tuned for both current and future prey position. Spatial efficiency (SPAEF), a measure of the similarity of two-dimensional filters^29^, is show on the y-axis of the central plot. A more positive SPAEF indicates that the matrices are more similar to each other; low values indicate orthogonality. Only significant neurons are shown; cells are sorted by spatial efficiency. **(D)** Sliding window analysis for future position encoding strength. Plot shows proportion of neurons significantly selective for future position at several possible future delays. This curve peaks at around 700-800 ms, which corresponds to the average prediction distance for all three subjects. **(E)** The distance between current prey position and future prey position at time t rises roughly linearly with time. This finding indicates that the peaks found in panel D are not likely to be an artifact of some unforeseen periodicity in the relative paths of the subject and prey.

We called our first model the *physics variable based prediction model* (PVBP). It assumes that subjects’ prediction derives from the the prey’s current position, velocity (i.e. both speed and direction), and acceleration (which includes both direction and magnitude of acceleration), as well as further derivatives, see Figure S5). For all three subjects, the best fitting τ is positive, indicating that they point the joystick towards the prey’s future position. For ease of interpretation, we translated τ into time units by calculating the distance between the current position and estimated position, then divided that quantity by the average velocity of the prey across the session. The results of this calculation indicate that subjects K, H and C pointed the joystick towards the position that the prey would occupy in an average of 800 ms, 767 ms, and 733 ms in the future, respectively. In the context of the task, these numbers are substantial: they reflect 18.78%, 14.42%, and 13.23% of the average trial duration for K, H, and C, respectively.

**Figure 5.**
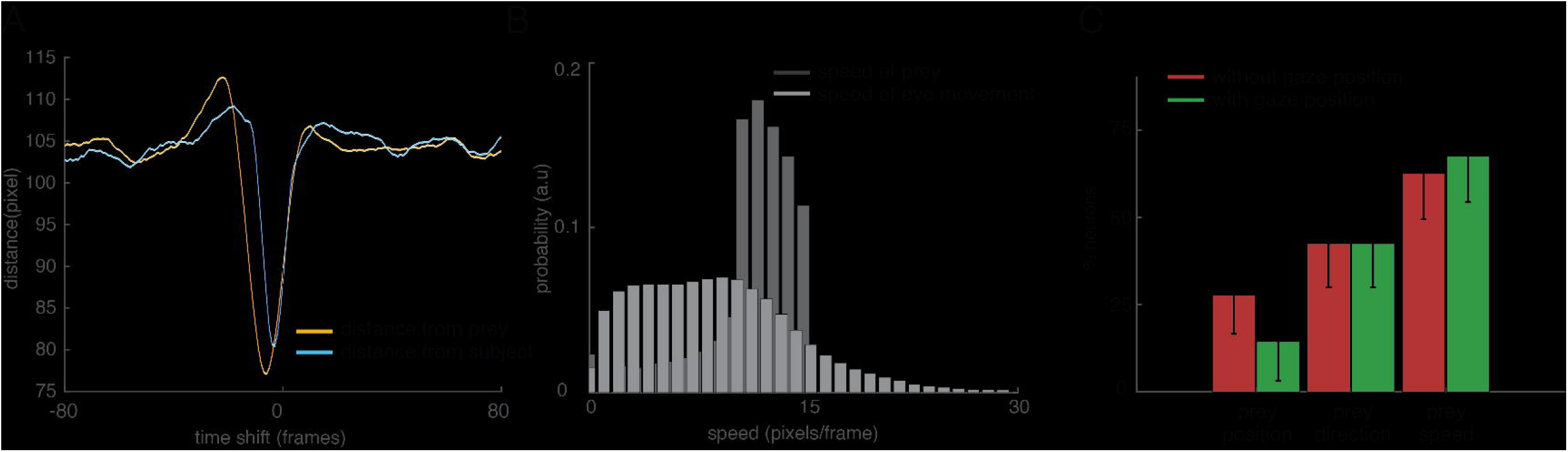
Analyses that control for potential gaze confounds. **(A)** The Euclidean distance between the eye position at t=0 and prey position (orange solid line) / self position (blue solid line). Error bar = SEM, and is the width of the lines shown. **(B)** Speed distribution of prey movement and smooth eye pursuit. **(C)** Proportion of neurons tuned for three key variables using the standard GLM described above and another version that assigns variance to eye position first. All three variables are still significant in the population when including gaze position.

To determine whether the positive prediction parameter τ is significantly greater than zero, we performed a bootstrap (randomization test making use of resampling with replacement) of heatmap slices from each segment (individual heatmap from 500 segments). This resampling was performed 500 times and resulting heatmaps were added. Then the τ and κ that best explain each segment (that is, the one resulting in the lowest cost) was selected in each resampling. We confirmed that the estimated value τ and κ are both greater than zero more than 99% of the time (i.e., p<0.01).

The distance into the future that our subjects predicted did not detectably depend on the the speed of the prey (linear regression between reward/speed and mean τ K: β =3.0316, p=0.1110; H: β=4.5798, p=0.1791; C: β=7.1007, p=0.0957; the term β refers to the regression coefficient for speed against neural activity). We next asked whether taking more complex paths (ones with more turns vs. more straight paths) affected prediction span. Prey path complexity (as measured by path curvature estimated by average angle method) affected prediction. Specifically, subjects predicted less far into the future when the prey path had more curves (K: β=−0.0687; H: β=−0.0567; C: β=−0.0898, p<0.0001 for each). Thus, subjects had the ability to β dynamically adjust their own prediction in light of changing circumstances.

### Alternative models do not predict trajectories as well as physics-based prediction

We next compared the physics-based model to two other models implementing different prediction algorithms (Figure 2B). First, the veridical prediction (VP) model assumes that the subjects will make perfect predictions that incorporate all game dynamics, including preys’ repulsion from the walls and the subject’s avatar. This means that a subject that makes a veridical prediction takes into account the effect his own movements will have on the prey’s strategy. Second, the cost contour map prediction (CCMP) model is the same as VP but excludes repulsion from the avatar, meaning that the subject’s prediction model for the prey would not take into account their own motion. We compared the performance of each model by computing the sum of squares error between the prediction trajectory and the observed trajectories over all time bins.

Using the Akaike Information Criterion (AIC), we found that the PVBP fit better than the other two models in our well-trained subjects (K: 7.529×10^6^, for subject K, second best was VP: 7.542×10^6^; H, PVBP: 8.923×10^6^; for subject H, second best was CMPP: 8.950×10^6^, Figure 2D). We fit each segment with distinct τ and κ parameters, and we fit these same two parameters for each of our three models. As a consequence, the comparison of models can be done directly without concern of potential bias toward any specific model. In other words, by fitting each of the three models subject to identical constraints, we ensured a fair comparison across models. For the less well-trained subject, C, the CCMP model explained trajectories most accurately (7.955×10^6^).

We speculated that one factor that may influence strategy is the speed of the prey. Indeed, we found that all three subjects used PVBP more frequently when the speed of the prey was faster (Figure S6). Note that this observed link between speed and the fit of the PVBP occurs even in our third (less fully trained) subject (p < 0.001, logistic regression, Figure S6). In any case, our model’s classification of strategies appears to be robust: the same results were obtained using a different method. Specifically, we fit all individual segments to the best model and computed the model that fit the most overall number of segments (Figure 2D and **Methods**).

**Figure 6.**
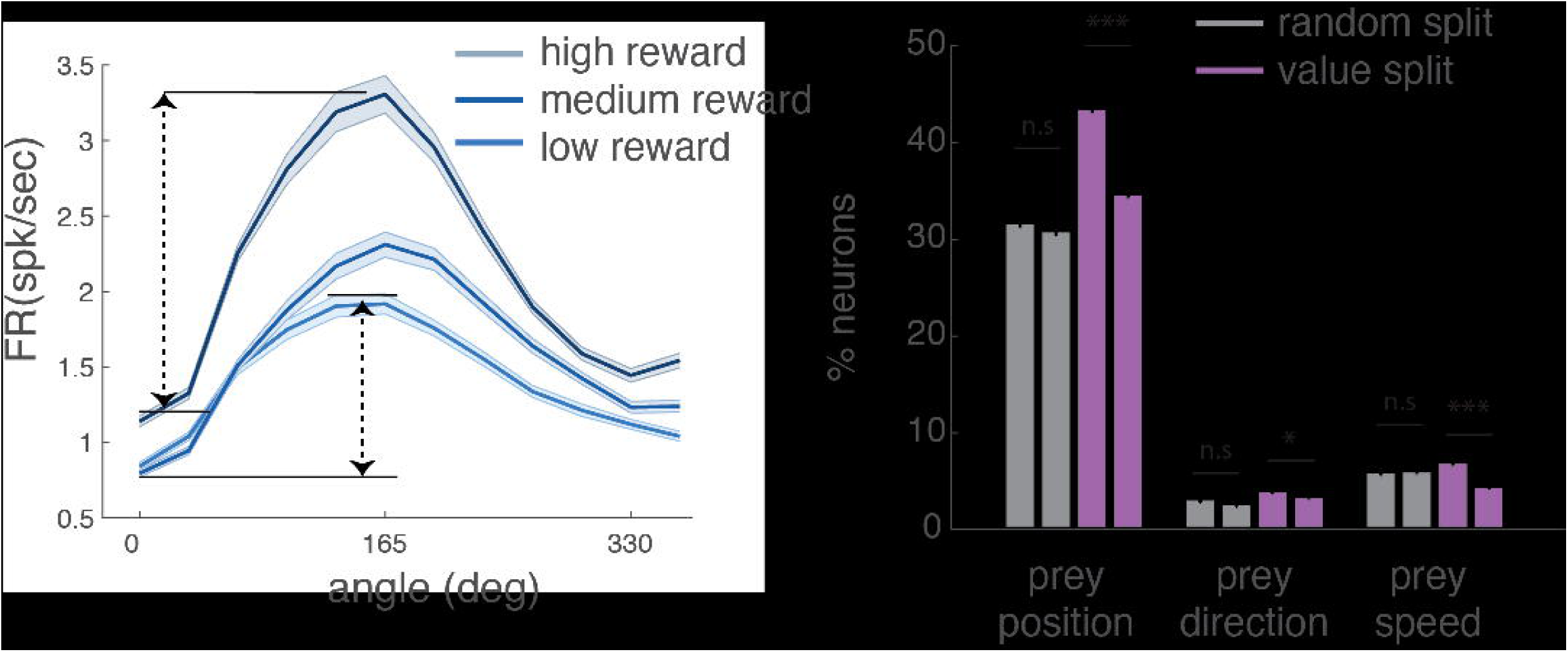
Modulatory effect of reward size on tuning for prey variables. **(A)** Responses of an example neuron selective for the angle between self and prey; changes in the reward size of prey (divided into three bins) appear to change the gain and not the offset of the neurons; that is, reward interacts multiplicatively with angle. **(B)** This pattern is also observed in the population. The proportion of neurons significantly tuned for prey variables (prey position, prey direction, and prey speed) when splitting data randomly (grey bar) or according to value of pursued prey (purple bar). The difference of value split was significant (p = 0.0221 for prey speed, and p < 0.001 for other prey variables).

Doing the fitting this way may seem excessively flexible. That is, using two times the number of segments might allow us to fit only noise. (Consider, for example, the case of fitting 9 data points with 9th-order polynomial curve). On the other hand, the extra freedom may allow us to better fit signal - or, of course, it may fit both noise and signal. The key question, then, is whether using a large number of parameters makes the fit better despite the possibility of fitting noise. To answer this question, we directly compared the two approaches (Figure S3).

Specifically, we compared a model assigning two parameters globally versus one applying two parameters for each one-second segment (i.e., Monkey K: 28,164; Monkey H: 35,308; Monkey C: 20,720 parameters, Figure S3). We then used AIC to compare models. We found that the second-by-second fitting resulted in lower AIC values, implying a better fit, than the 2-parameter counterpart. Specifically, in this figure, for all individual subjects, the change in AIC (AIC for global parameter model minus AIC for second-by-second model) was positive - implying the model fit by second-by-second model explains the data better than the counterpart - for the best physics-based model explaining the subject’s behavior.

Overall, the model comparison results showed that subjects predict the upcoming position of the prey using Newtonian physics but ignore the walls and their own influence on the prey. That is, subjects use a simplified approximation of the structure of the game to make future predictions; presumably this simplified one is sufficient to generate good predictions with lower mental effort costs. Indeed, the correlation between speed of prey and subjects’ reliance on physics based prediction (a result confirmed with two different analytical approaches) suggests that prediction might have a computational cost.

### Prediction-related information encoded in dorsal anterior cingulate cortex

Based on its role as a nexus for motivational, cognitive, and motor information ^18, 23^, and its demonstrated role in human prospection ^11, 25^, we hypothesized that dACC would be critical for predictive pursuit (Figure 3A). We fit a statistically unbiased Linear-Nonlinear Generalized Linear model (GLM ^26–28^) to responses of 150 well-isolated dACC neurons (K: n=31; H: n=119). For this analysis, we focused on the entire trial period rather than pre-selecting epochs.

Position, velocity, and acceleration of the prey were all encoded by significant proportions of neurons (Figure 3; position: 62.00%, n=93/150; speed: 35.33%, n=53/150; 36.67%; direction: n=55/150, acceleration: 24.67%, n=37/150, p<0.01 in all cases, two-way binomial test). The model fit shown in magenta is the shape of reconstructed filter (examples, Figure 3C, D). According to the GLM, jerk, the derivative of acceleration, is not encoded (see Figure S5). Jerk also did not measurably affect the subject’s neural responses (it only modulated 2.00% of cells, n=3/150, p=0.1288). Together, these results indicate that dACC ensembles carry the major raw ingredients that our subjects use to predict prey positions.

We wondered whether ostensible coding for prey variables could be the byproduct of coding for self-position, since self-position and prey position do tend to be somewhat correlated. We therefore repeated our GLM analyses but included self-position, self-direction, and self-speed as explanatory factors and considered variance explained by prey parameters only after accounting for these variables. Doing this, the proportion of neurons selective for the prey’s position information remained significant (position: 65.45%; p < 0.01, two-way binomial test), as did neurons selective for prey speed (18.56%; p<0.01), and direction (10.78%, p=0.021).

### Neurons in dACC encode future position

We next asked whether dACC neurons encode the future position of the prey. For each neuron, we refit the GLM using an additional parameter, the position of the prey at time t in the future. We selected the time t (t=833ms) that was most similar to the value of τ resulting from our generative model, that is, the one indicating the most likely time span of prediction (733, 766, and 800 for the three subjects, respectively) subject to the additional constraint of being a multiple of 166.67 (i.e. 10 frames). Note that although this value was chosen in advance, it aligns with the empirically derived measure of peak future position coding (Figure 4D, see below).

Our analysis approach deals with the problem of correlation between the set of current Newtonian variables (including current position) and future position by assigning all explanatory power to the set of current variables first, and only counting as significant any additional variance explained by future position (see **Methods**). Despite this conservative criterion, we found that responses of 24.67% of dACC neurons are selective for the prey’s future position at time t (n=37/150).

Visual inspection of the neurons’ filters shows that their selectivity is complex (examples are shown in Figure 4C). That is, they are positionally tuned, but, unlike place cells, have non-point-like shapes. They contain multiple peaks. They do not appear to be smooth gradients. Instead, they appeared to be heterogeneously spatially tuned. In this manner, they resemble recently identified non-grid-like space-selective cells in entorhinal cortex ^26^. Notably, conventional methods for detecting place/grid-like cells will greatly underestimate the proportion of such tuning.

We next asked how *strongly* dACC neurons encode the future position of the prey. We calculated the proportion of log likelihood increase (LLi) between the current position model and the current plus future model (Figure 4A). Our neurons showed a wide range in marginal variance explained. On average, adding the future position term improved variance explained by 6.89% (the mean of this proportion is significantly different from zero, p<0.001, Wilcoxon sign-rank test, Figure 4A **inset**).

We then asked whether these newly discovered *future position cells* constitute a separate class of neurons from the cells that tracked the current position of the prey. To do this, we computed the explanatory variance accounted for by future position (variance explained by the combined model minus variance explained by current position) and current position, as defined by log likelihood improvement (LLi) in fitting. We found a positive correlation between these variables (Figure 4B), indicating current and future position were multiplexed in the same population of cells (r = 0.7394, p < 0.001, cf. ^29^).

To quantify the difference between current and future position coding, we fit separate models: one incorporated current position plus current Newtonian variables; the other was the same but used future position (assuming t=833 ms) instead of current position. For the 36 neurons with significant tuning for both current and future position, we calculated the similarity between the filters, using a technique known as *spatial efficiency,* SPAEF ^30^ (Figure 4C). A zero SPAEF indicates orthogonal filters; positive SPAEF indicates similar filters; negative SPAEF indicates anticorrelated filters. Although the mean of the spatial efficency for our neurons was positive, it was not significantly so, and spanned a large range of values from negative to positive (mean of population spatial efficiency=0.0440, Wilcoxon sign-rank test, p=0.3790).

Finally, we assessed future encoding by examining the accuracy of model fitting to each of several possible future times, ranging from 0 to 1333 ms in the future. We ran a type of sliding window analysis that involved sampling one frame (16.67 ms) every ten frames (166.67 ms) and ignoring the intervening nine frames. We found that the value of 833 ms fit the largest number of neurons. (Values around it fit many neurons too). Specifically, the plurality, 24.67% of neurons, were tuned for prey position at 833 ms (Figure 4D). The roughly equivalent value of the neural and the behaviorally fit prospective distance (733 and 800 ms for those two subjects) suggests that these neurons encode the future position of prey on the same approximate timescale as the subject actively predicts.

We considered the possibility that this peak at 833 ms was due to some unanticipated correlation between positions in the future and at the present. If this were so, then the average distance of the self and/or prey would show a local minimum at a point in the future corresponding to the peak. However, we did not see this. On the contrary, we found that the distance increases monotonically for both subjects (Figure 4E).

### State information is not confounded with gaze information

Activity in dACC is selective for saccadic direction, and may therefore also correlate with gaze direction (although this has not, to our knowledge, been shown ^31^). Consequently, it is possible that our spatial kernels may reflect not task state but gaze information. Specifically, what appears to be tuning for future position may instead be attributable to the fact that monkeys looked towards the predicted future prey position. We tested this possibility by calculating the Euclidean distance between eye position and prey position in a range from −80 to +80 frames (Figure 5A). The distance between eye and prey position was the closest at −5 frames (77.09 pixels), indicating that eye position *lagged* prey position. Thus, if gaze direction were a major confound, it would show up as increased selectivity for past positions, not prediction of future positions. Likewise, the chance that prey velocity encoding is a by-product of eye velocity encoding was belied by the stark differences between gaze speed and prey speed (p <0.001, Wilcoxon sign rank test, but also clear from visual inspection of Figure 5B). Finally, we repeated our GLM analyses (see above) but included eye position (only for the one subject from which we collected gaze data). We found that that the number of tuned neurons for the prey did not substantially change; that is, that adding in gaze position as a regressor did not qualitatively change our results (Figure 5C).

### Encoding of reward and reward proximity in dACC

Research based on conventional choice tasks indicates that dACC neurons track values of potential rewards ^32^. We next asked how dACC encodes anticipated rewards in our more complex task. We found that, averaging over all other variables, the value of the pursued reward modulates activity of 8.67% of neurons (using a simple linear regression of firing rate against value; this proportion is greater than chance, p=0.038, one-way binomial test). Note that this analysis ignores the potential encoding of prey speed, which is perfectly correlated with static reward in our task design. We hypothesized that reward/speed would be encoded in a modulatory manner ^33^, that is, that the pursued reward/speed would alter the shape of the tuning for other task variables, rather than be multiplexed (Figure 6A). To test this hypothesis, we split our dataset by reward size and, as a control, split it randomly. We found that for several variables (prey position, prey direction, and prey speed), value splits produced greater differences than random ones (purple bar, p = 0.0221 for prey speed, and p < 0.001 for other prey variables, Figure 6B). This result indicates that the reward information encoded in dACC interacts mathematically with encoding of other variables. In other words, selectivity is mixed.

A good deal of research suggests that dACC neurons also signal the approach in time of impending rewards ^34–36^, even in continuous tasks ^37, 38^. We thus asked whether it does so here. We repeated our GLM, including relative (self-to-prey) distance as an explanatory variable. We found that 38.67% of neurons (n=58/150) were tuned for self-prey distance. Interestingly, this relationship is heterogeneous - of these 58 neurons, 31.03% (n=18/58) showed a positive slope and 18.97 % (n=11/58) showed a negative slope. This bias is not itself significant (p=0.2649 for rise and fall bias, n = 18/29; p = 1.000 for monotonic bias, n = 29/59, binomial test in all cases). This result indicates that while dACC neurons do track the approach to reward, they do not show an overall rise or fall in activity as they do so.

## DISCUSSION

Pursuit is an important element of the behavioral repertoire of many foragers ^2, 6^. The algorithmic bases of pursuit have recently attracted the interest of scholars in ecology, engineering, psychology and other disciplines ^4, 7, 39–44^. Nonetheless, we know very little about how pursuit decisions occur in real time, and we know even less about their neuronal underpinnings. Here, we examined how macaques pursue virtual prey in a continuous, time-varying task. We developed a generative model based on a large dataset. The result from this model suggests that our subjects follow a predictive strategy. That is, instead of pointing towards the position of the prey, they extrapolate the future positions of prey and use this prediction to adjust their heading. This strategy is more efficient (yields more reward per unit time) but may be more computationally demanding than a simpler one that would involve pointing at and tracking the current position of the prey. These results demonstrate that pursuing animals can adopt complex future-predicting strategies that improve performance.

We found that dACC neurons track the elemental physical variables our subjects use to predict the future and explicitly encode the prediction. Specifically, we found that firing rate responses of neurons in dACC encode three Newtonian variables (position, velocity, and acceleration) that our subjects used to track the prey and predict future prey positions. The same neurons carry an additional representation of the future position of the prey that is multiplexed with the Newtonian variables rather than maintained in a separate pool of specialized neurons. Both representations make use of a two-dimensional response field, akin to place fields in hippocampus, but not localized to a single position. Specifically, spatial representation in dACC is qualitatively similar to place representations of non-grid cells in entorhinal cortex ^26^. It is notable that dACC uses partially distinct spatial tuning functions to track the present and future positions of the prey, thus in principle allowing unambiguous decoding for a given population response.

Our work is directly inspired by important studies identifying mechanisms underlying pursuit in other animals ^39, 40, 45^. Our work goes beyond these studies by developing a generative model, that is, a model that seeks to understand how the data are generated ^46^. One benefit of the generative model is that it lets us probe how the decision is made at every time step and make guesses about the underlying mental process leading to decision. The generative model in turn is vital for extending our understanding of mechanism to the neuronal level.

This model allows us to generate results that provide novel insight into the role of dACC in cognition. First, our results emphasize the core role of dACC in prediction, a role that is central to other theories, albeit not ones that directly involve pursuit ^11, 17, 20, 21, 47, 48^. One recent study is particularly relevant to these results ^20^. The authors examined hemodynamic activity in human dACC during a complex decision-making task in which subjects had to track previous rewards and use a reinforcement learning-like mechanism to formulate a future prediction and make the best choice. They found that dACC tracks multiple variables, but was particularly selective for long-term estimates of expected prediction errors. These results highlight the key role of dACC in prediction in general and suggest its role is conserved across species (see also ^17^). Second, our findings highlight the importance of dACC to navigation. While studies of navigation typically focus on the medial temporal lobe, a growing body of work has begun to explore the role of cingulate cortex, which receives direct projections from medial temporal regions ^25, 49^.

There are several important limitations to the present work. First, and most obviously, our subjects were not performing a truly naturalistic task; they were performing a laboratory task that required specialized training. Future studies will be needed to ascertain whether these results relate to natural pursuit contexts that are ostensibly similar, such as pursuit of insects in the peripersonal space ^50, 51^. Second, and relatedly, the task space we used was greatly constrained - both agents were restricted to a small rectangular space and had strict speed limits. Subjects had full information about the position of the prey at all times. To understand prediction more fully, it will be critical to extend to contexts in which some information is hidden.

Traditional laboratory tasks that study topics of interest to cognitive neuroscience - decision-making and executive control - have discrete steps and force the brain to adjust to that structure ^52, 53^. One reason we developed the prey pursuit task is that it embeds those cognitive processes in a continuous time-varying task. Doing so allows us to study one of the brain’s greatest strengths - its ability to adjust and change its mind on the fly as new evidence comes in ^53–57^, and to incorporate that into future plans.

## Acknowledgements

We thank Alex Thomé for his critical role in designing the task, for devising the training protocols, and for developing our joysticks. We thank Marc Mancarella for his critical help with joystick training. We appreciate invaluable help from Marc Schieber, Adam Rouse, and Sarah Heilbronner.

## Author Contributions

SBMY and BYH conceptualized and designed the experiment. SBMY collected the data. SBMY, and STP developed the behavioral model, SBMY, JCT, and BYH developed the physiological model and analyzed the data. SBMY and BYH wrote the manuscript.

## Supplementary Material

### Material and Methods

#### Subjects

All animal procedures were approved by the University Committee on Animal Resources at the University of Rochester and/or the University of Minnesota and were designed and conducted in compliance with the Public Health Service’s Guide for the Care and Use of Animals. Three male rhesus macaques (*Macaca mulatta*) served as subjects for the behavior; two of them also served as subjects for the physiology. Subjects had never previously been exposed to decision-making tasks in which they could use a joystick to pursue a moving prey. Previous training history for these subjects included two types of foraging tasks ^37, 57^, intertemporal choice tasks ^59^, several types of gambling tasks ^60–62^, attentional tasks (similar to those in ref ^63^), and two types of reward-based decision tasks ^64, 65^.

#### Experimental Apparatus

The joystick was a modified version of commercially available joysticks with a built-in potentiometer (Logitech Extreme Pro 3D). The control bar was removed and replaced with a control stick (a 15 cm plastic dowel) topped with a 3 cm diameter plastic sphere designed to be easy for macaques to manipulate. The joystick position was read out by a custom coded program in Matlab running on the stimulus-control computer. The joystick was controlled by an algorithm that detected the positional change of the joystick and limited the maximum pixel movement to within 23 pixels in 16.67 ms.

#### Task Design

At the beginning of each trial, two shapes appeared on a gray computer monitor placed directly in front of the subject. The yellow (subject K) or purple (subjects H and C) circle (15-pixel diameter) represented the subject. Subject position was determined by the joystick and was limited by the screen boundaries. A square shape (30 pixel length) represented the prey. The movement of the prey was determined by a simple AI (see below). Each trial ended with either the successful capture of the prey or after 20 seconds, whichever came first. Successful capture was defined as any spatial overlap between the avatar circle and the prey square. Capture resulted in immediate juice reward; juice amount corresponded to prey color: orange (0.3 mL), blue (0.4 mL), green (0.5 mL), violet (0.6 mL), and cyan (0.7 mL).

The path of the prey was generated interactively using A-star pathfounding methods, which are commonly used in video gaming ^66^. For every frame (16.67 ms), we computed the cost of 15 possible future positions the prey could move to in the next time-step. These 15 positions were spaced equally on the circumference of a circle centered on the prey’s current position, with radius equal to the maximum distance the prey could travel within one time-step. The cost in turn was based on two factors: the position in the field and the position of the subject’s avatar. The field that the prey moved in had a built-in bias for cost, which made the prey more likely to move towards the center (Figure 1B). The cost due to distance from the subject’s avatar was transformed using a sigmoidal function: the cost became zero beyond a certain distance so that the prey did not move, and it became greater as distance from the subject’s avatar decreased. Eventually, the costs from these 15 positions were calculated and the position with the lowest cost was selected for the next movement. If the next movement was beyond the screen range (1920×1080 resolution), then the position with the second lowest cost was selected, and so on. The maximum speed of the subject was 23 pixels per frame (and each frame was 16.67 ms). The maximum and minimum speeds of the prey varied across subjects and were set by the experimenter to obtain a large number of trials (Figure 1). Specifically, speeds were selected so that subjects could capture prey on <85% of trials; these values were modified using a staircase method. If subjects missed the prey three times consecutively, then the speed of the prey was reduced. Once the subject intercepted the prey in a trial where the staircase method was used, then the selection of prey speed was randomized again. To ensure sufficient time of pursuit, the minimum distance between the initial position of each subject avatar and prey was 400 pixels.

#### Training Level Estimation

Three subjects were trained for the same amount of time (8 weeks). As training progressed, each subject was exposed to a progressively more difficult (faster) suite of prey, up to a fixed maximum. Subject K and subject H reached a similar range for maximum speed of prey during the training period (K:15 pixels per frame; H: 14 pixel per frame). However, subject C only attained a maximum speed of 8 pixels per frame (Figure S6). It is for this reason we refer to him as the less well-trained subject.

#### Behavioral Model

To fit each subject’s movement, each trial was divided into 1 second-long segments. Each segment included 61 data points (because we used 16.67 ms resolution).

We modeled these trajectories using a single prediction and a single force parameter for the entire trial, as a simplifying assumption. Nonetheless, it is reasonable to assume that throughout a long, 20-second period, there would be active adjustment of prediction and force. Actual comparison by AIC supported our intuition, and we used segment as the unit of analysis throughout (values of ‘AIC of segment/AIC of trial’ was 0.9328, 0.9214, 0.9227, for subjects K, H, and C (or whatever) respectively.

Overall, the position of the subject was generated according to the following:

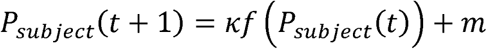

where P_subject_(t) is position of the subject at time t, m is the inertia of subject as calculated from the joystick, and κ is the force parameter. The vector □*f*(P_subject_(t)) was then summed with the inertia m that was defined as following:

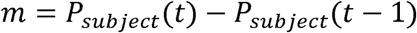

P_prey_(t) indicates the position of the prey at time t. The function with respect to subject position at time t was defined as:

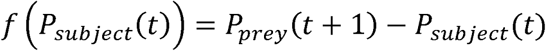

Then the position of the prey at time t+1 was:

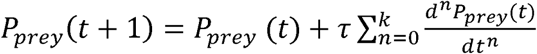

Where the n indicates the order of derivation with respect to the time. Thus, n=1 indicates velocity, and n = 2 indicates acceleration.

The Physics Variable-Based Prediction (PVBP) model incorporates one previous time step to predict the prey’s next position. This approach is similar to a Kalman filter ^67^. The other two models we tested do not utilize any past information. The model assuming prediction using the cost contour map (CCMP model) considers only the lowest cost location at the next time step. The model assuming veridical prediction (VP) automatically finds the exact position of the prey at the next time step. Once the prey’s position on the next time step is predicted, the model computes how far this predicted position is from the agent’s current position. A prediction value of 1 indicates that the future position will be as far as from the agent’s current position as the prey’s current position. The best-fitting parameter pairs were determined by performing a grid search across the ranges of both parameters.

During this search, we tested the range of the prediction parameter between −400 to 400 subjects H and C, and −200 to 200 for subject K. (Units for this range correspond to the distance the prey moved in the previous timestep). Subjects H and C had a larger range, because over 5% of their trajectories resulted either in −200 or 200 in prediction parameter value. Representative parameters for explaining each segment were selected based on the value of the sum of squared error between the actual segment and the segment generated by the model.

#### Significance Testing

To determine whether the positive prediction parameter was significantly greater than zero, we performed a bootstrap of heatmap slices from each segment. This resampling was performed 500 times, and selected heatmaps were added. Then, the parameter set resulting in the lowest cost was selected in each resampling.

#### Model Evaluation

To evaluate model performance and compare among models, we computed the Akaike Information Criteria (AIC) using the likelihood of each model (Figure 2, and Figures S4 and S5). We first calculated the mean and variance of all the sum-of-squared errors across trajectories. Then we estimated the likelihood assuming a normal distribution centered on the mean of the sum-of-squared errors with a variance equivalent to the variance of the sum-of-squared errors across all trajectories. To validate whether subjects used a single prediction and force across the all the trials or adaptively changed their prediction method, we compared the AIC value between cases where the parameter pair varied across all trajectories, using only the single best parameter pair.

#### Electrophysiological recording

One subject (H) was implanted with multiple floating microelectrode arrays (FMAs, Microprobes for Life Sciences, Gaithersburg, MD) in the dorsal anterior cingulate cortex (dACC). This is the region that we define as Area 24 ^18^ and that corresponds to dACC in most other primate studies, including those from our lab ^37, 60, 68^. Each FMA had 32 electrodes (impedance 0.5 MOhm, 70% Pt, 30% Ir) of various lengths to reach multiple layers within dACC. Neurons from subject K were recorded with laminar V-probes (Plexon, Inc, Dallas, TX) that had 24 contact points with 150 μm inter-contact distance. Continuous, wideband neural signals were amplified, digitized at 40 kHz and stored using the Grapevine Data Acquisition System (Ripple, Inc., Salt Lake City, UT). Spike sorting was done manually offline (Plexon Offline Sorter). Spike sorting was performed blind to any experimental conditions to avoid bias.

#### Details of LN model

To test the selectivity of neurons for various experimental variables, we constructed Generalized Linear Models with navigational variables (GLM ^26, 28^). The GLM models estimated the spike rate (r_i_) of one neuron during time bin t as an exponential function of the weighted sum of the relevant value of each variable at time t, which the weights are determined by set of coefficients (w_i_). The estimated firing rates from the GLM models can be expressed as:

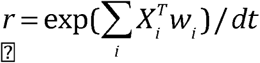

Where r denotes a vector of firing rates for one neuron over T time points across the session, and i indexes the variables of interest, e.g. position of avatar on screen. The vector of firing rates over T time points provides the benefit for modeling the neural activity without specific time-locking to behavioral event. X_i_ is a matrix in which each column represents a set of “state variables” of the animal (e.g. one of twelve speeds, determined by post-hoc binning) obtained from binning the continuous variable so that all the columns for a particular row are 0, except for one column. Unlike conventional tuning curve analysis, GLM analysis does not assume the parametric shape of the tuning curve *a priori*. Instead, the weights, which define the shape of tuning for each neuron, were optimized by maximizing the Poisson log-likelihood of the observed spike train given the model-expected spike number, with additional regularization for the smoothness of parameters in a continuous variable, and a lasso regularization for parameters in a discrete variable. Position parameters were smoothed across rows and columns separately. The regularization hyperparameter was chosen by maximizing the cross-validation log-likelihood based on several randomly selected neurons. The unconstrianed optimization with gradient and Hessian was performed (MATLAB fminunc function). Model performance of each neuron was quantified by the log-likelihood of held out data under the model. This cross-validation procedure was repeated 10 times (10-fold cross-validation), and overfitting was penalized. Through multiple levels of penalties, we can compare performance of models with varying complexity.

#### Forward model selection

Model selection was based on the cross-validated log-likelihood value for each model. We first fit *n* models with a single variable, where n is the total number of variables. The best single model was determined by the largest increase in spike-normalized log-likelihood from the null model (i.e., the model with a single parameter representing the mean firing rate). Then, additional variables (n-1 in total) were added to the best single variable model. The best two-variable model was preferred over the single variable model only if it significantly improved the cross-validation log-likelihood (Wilcoxon Signed Rank Test, α = 0.05). Likewise, the procedure was continued for the three-variable model and beyond if adding more variables significantly improved model performance, and the best, simplest model was selected. The cell was categorized as not tuned to any of the variables considered if the log-likelihood increase was not significantly higher than baseline, which was mean firing rate of fitted neurons across the session.

#### Future position models

We examined effect of future position by fitting a GLM having ‘future position’ and ‘current position’ together as the input variable. Then we compared to the GLM model with only current position. Difference between the two models was evidence that additional variance was explained by including future position.

#### Comparison between current and future position filters

For this purpose, we constructed two GLMs: one with current position and current Newtonian variables (velocity and acceleration), and another with future position and current Newtonian variables. Then we selected the neurons that showed significant tuning for both models. To compare the similarity between two positional filters, we used the SPAtial EFficiency metric (SPAEF) that prior literature suggests to be more robust than the 2D spatial correlation ^30^. It quantifies the similarity between two maps:

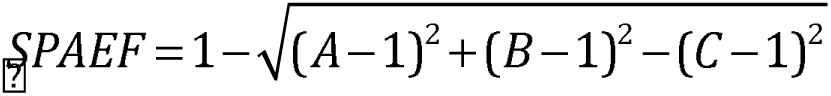

A is the Pearson correlation between two maps, B is the ratio between the coefficients of variation for each map, and C is the activity similarity measured by histogram profiles. Values near −1 indicate anticorrelated maps (one tends to be high when the other is low); 0 indicates uncorrelated maps; 1 indicates perfect matching between the two.

#### Velocity Dependent Physics Variable-Based Model (PVMP) Prediction Bias

We examined whether PVBP is preferred when the velocity of prey is high (Figure S6). We first obtained the average velocity of the prey at each segment, and then categorized each segment as belonging to either the physics or non-physics variable-based prediction based on which fit result was best. With the prey velocity and segment category, we performed logistic regression with velocity as a predictor and category as the dependent variable (glmfit in MATLAB).

#### Data availability

The datasets generated during the current study are available on the Hayden lab website, http://www.haydenlab.com/, or from the authors on reasonable request. The code generated to perform the analyses for the current study is available from the corresponding author.

## Supplementary Figure Captions

**Figure S1.**
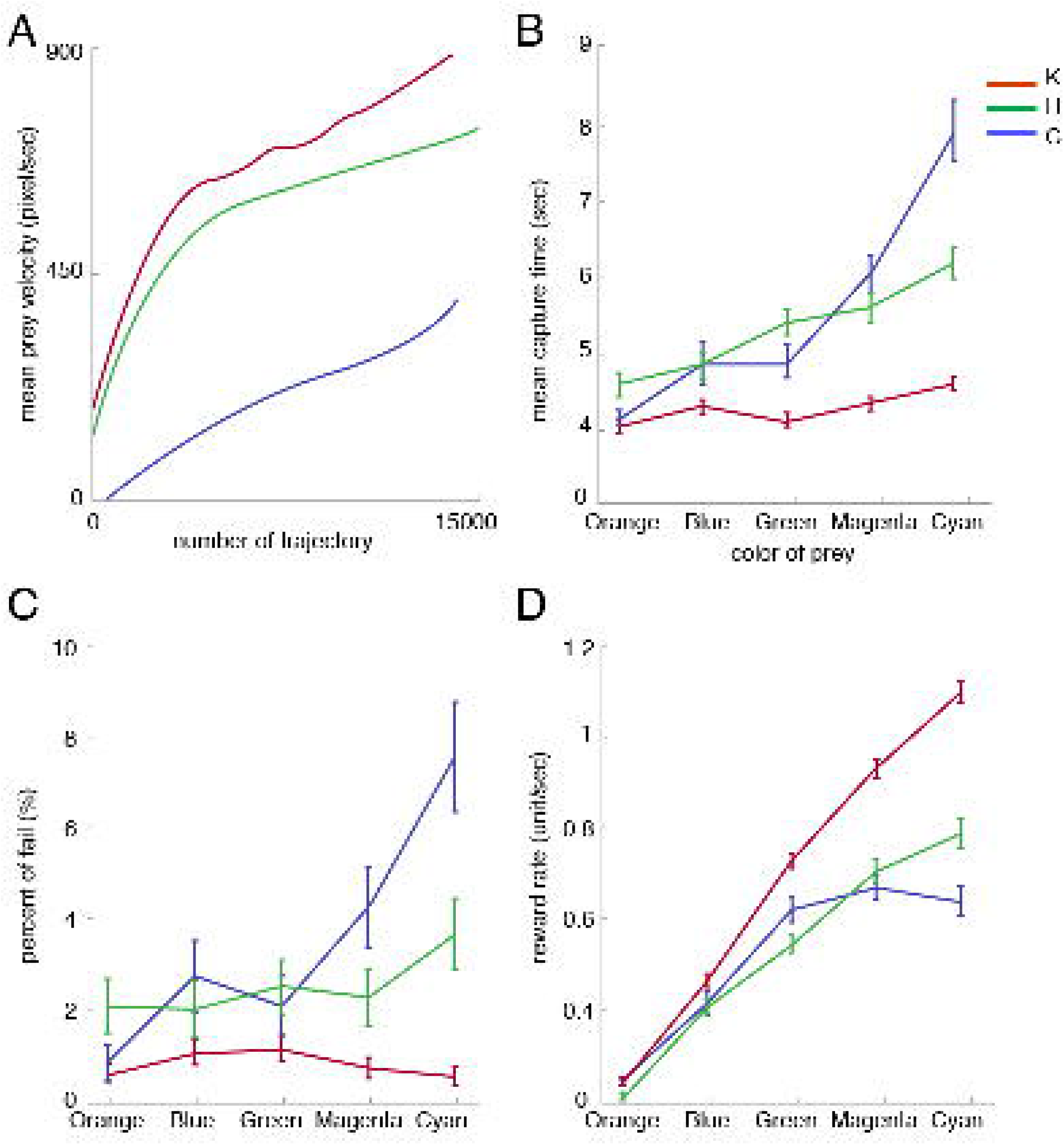
Subjects’ behavior varies according to prey speed/reward. (**A**) Mean prey velocity in each segment plotted separately for each subject. Pursuit result differs according to color (equivalent to maximum speed) of prey. The maximum speed of prey increases from orange (slowest with smallest reward) to cyan (fastest with largest reward). As maximum speed increases, the mean capturing time (**B**) and percent of failed trials increases (**C**). However, reward rate also increases, since the amount of reward is larger for faster prey (**D**). Errorbars are the standard error of the mean, obtained by bootstrapping (1000 bootstraps).

**Figure S2.**
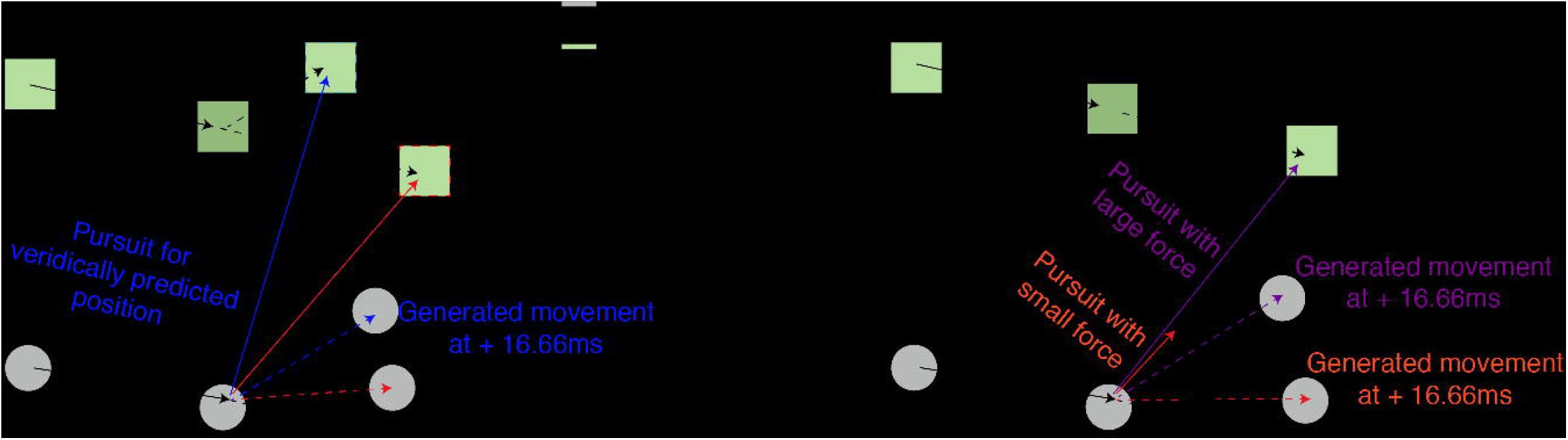
Different prediction strategies and influence of different amounts of inserted force. (A) The strategic difference between physics variable based prediction (PVBP, red lines) and veridical prediction (VP, blue lines). This generates different predictive points. (B) Effect of inserted force, shown between small (pink) and large (purple) forces. Vector-summation with inertia yields different outcomes for different force conditions.

**Figure S3.**
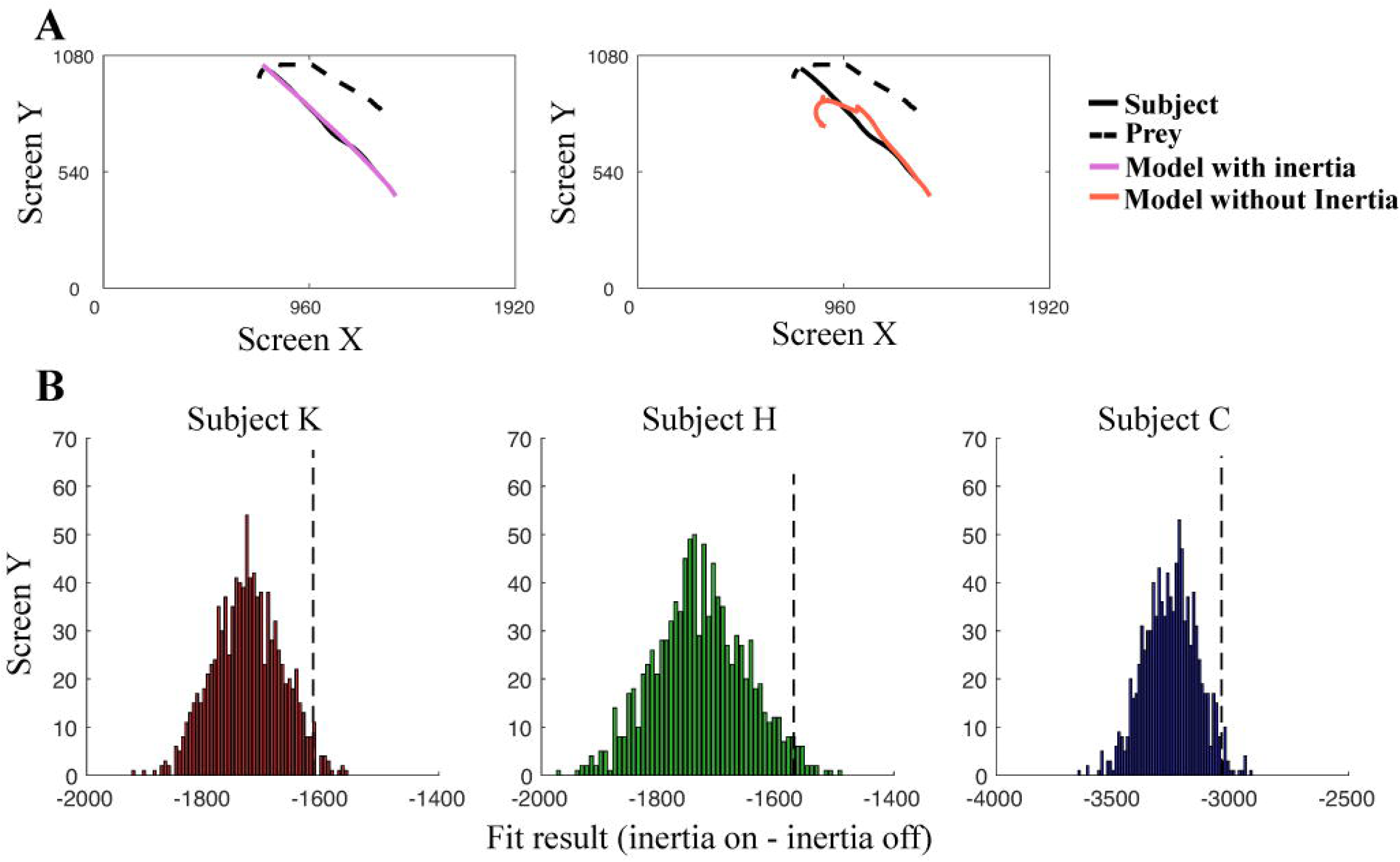
Dynamic changes of parameter sets at each segment explain each subject’s segments better than identical single parameter set across all the trajectories. AIC comparison between the case of the single parameter set across all the sessions (case 1) or adaptively changing parameter set at each segment (case 2). Delta AIC indicates the difference between the cases (case 1 - case 2), and a positive value indicates adaptively changing the strategy explains subject’s segment better, even if there is a penalty for having more parameters. Each column shows an individual subject’s result.

**Figure S4.**
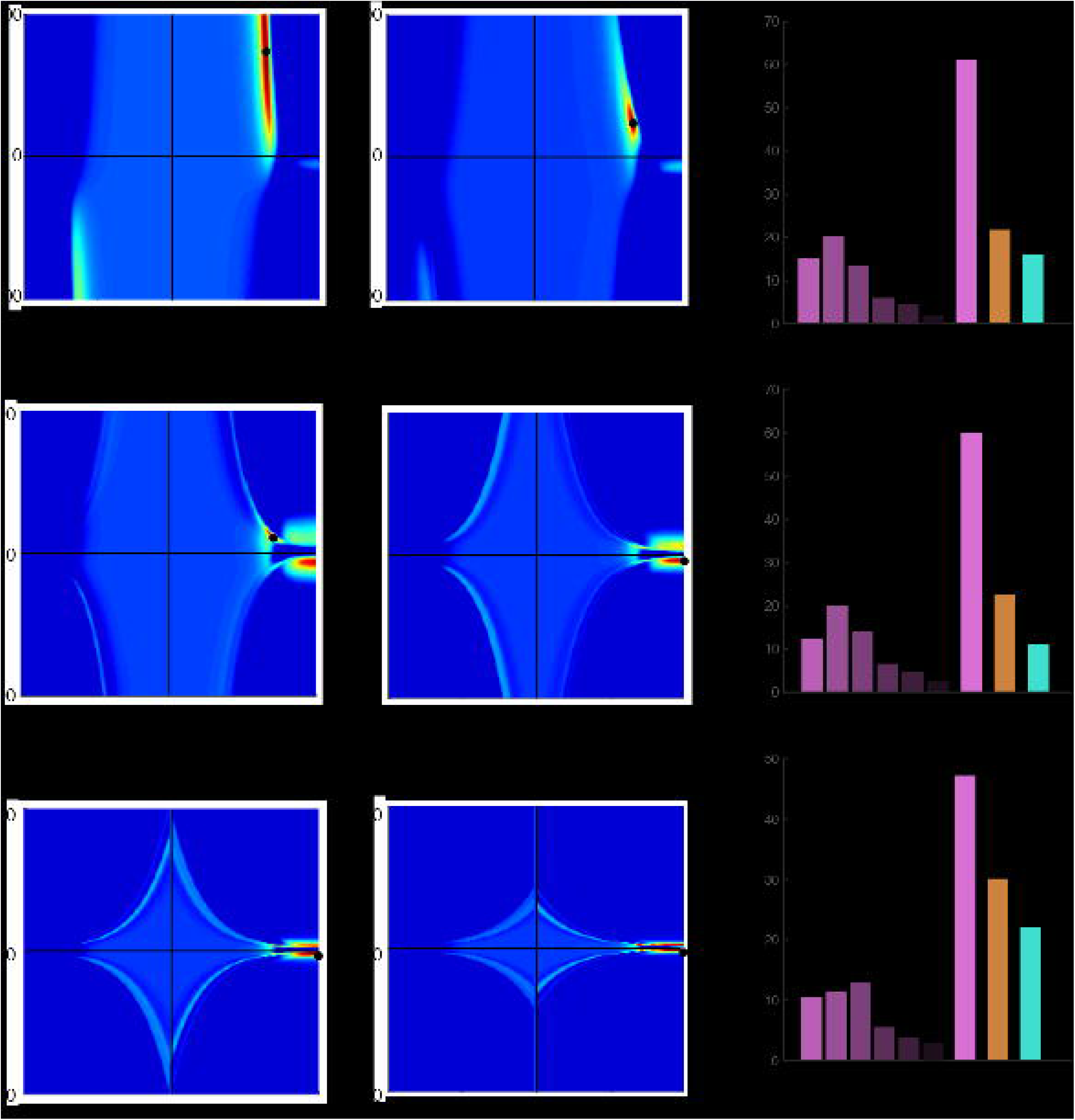
Including an inertia term improves model performance. (**A**) Model segment comparison between models with and without inertia. (**B**) Histogram results suggest that incorporating an inertia component to the model leads to a better fit of the data (mean of sum-of-squared error difference below zero at x-axis). 95% of data fall to the right of the black, dashed line. Bootstrapping of difference in performance between the model with and without inertia was performed in randomly sampled trajectories (number of resamples: 1000; randomly selected trajectories: 2000).

**Figure S5.**
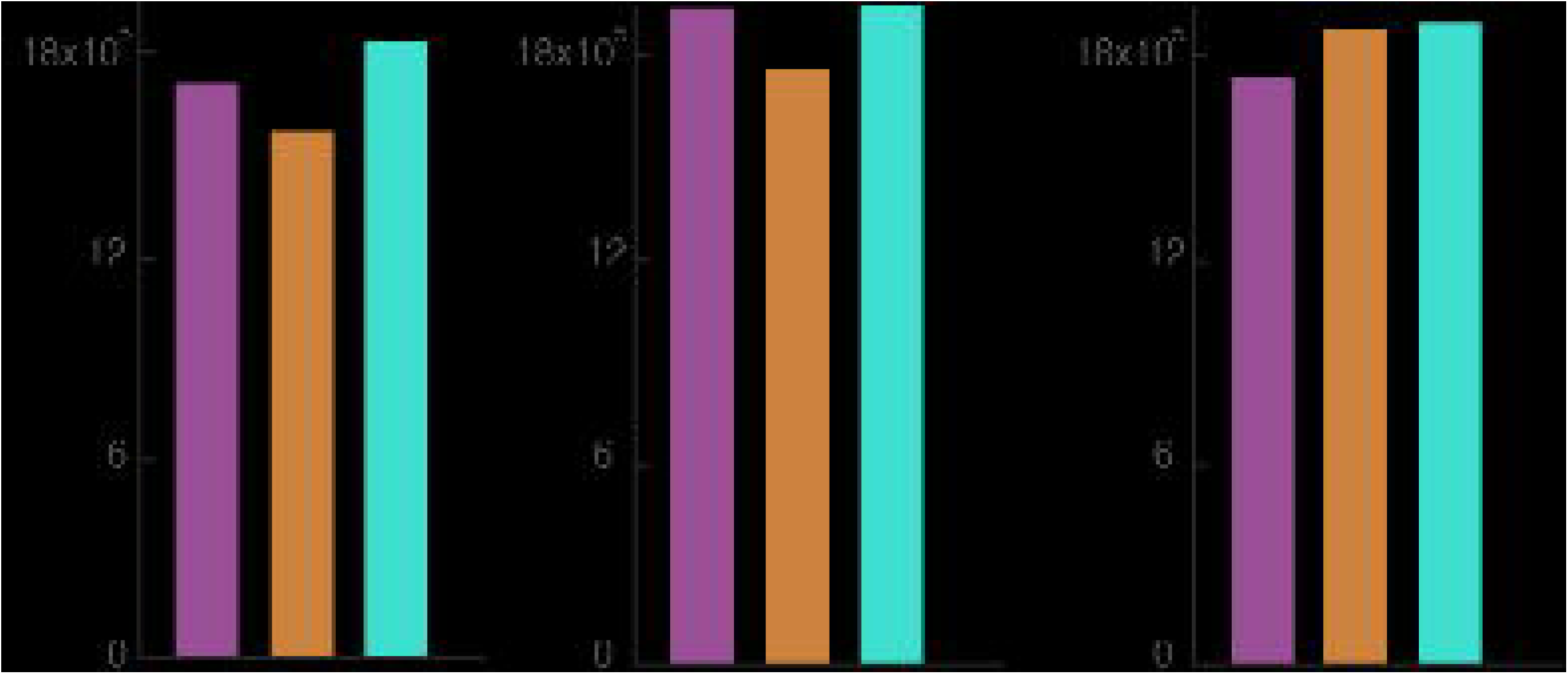
Additional terms after acceleration don’t improve model performance. (A) Each heatmap indicates the addition of more physical derivatives of position. The black circle indicates the best parameter set for the model. (B) Summary bar graph. Physics include within-physics prediction model comparison (from velocity to pop, the 6th derivative).

**Figure S6.**
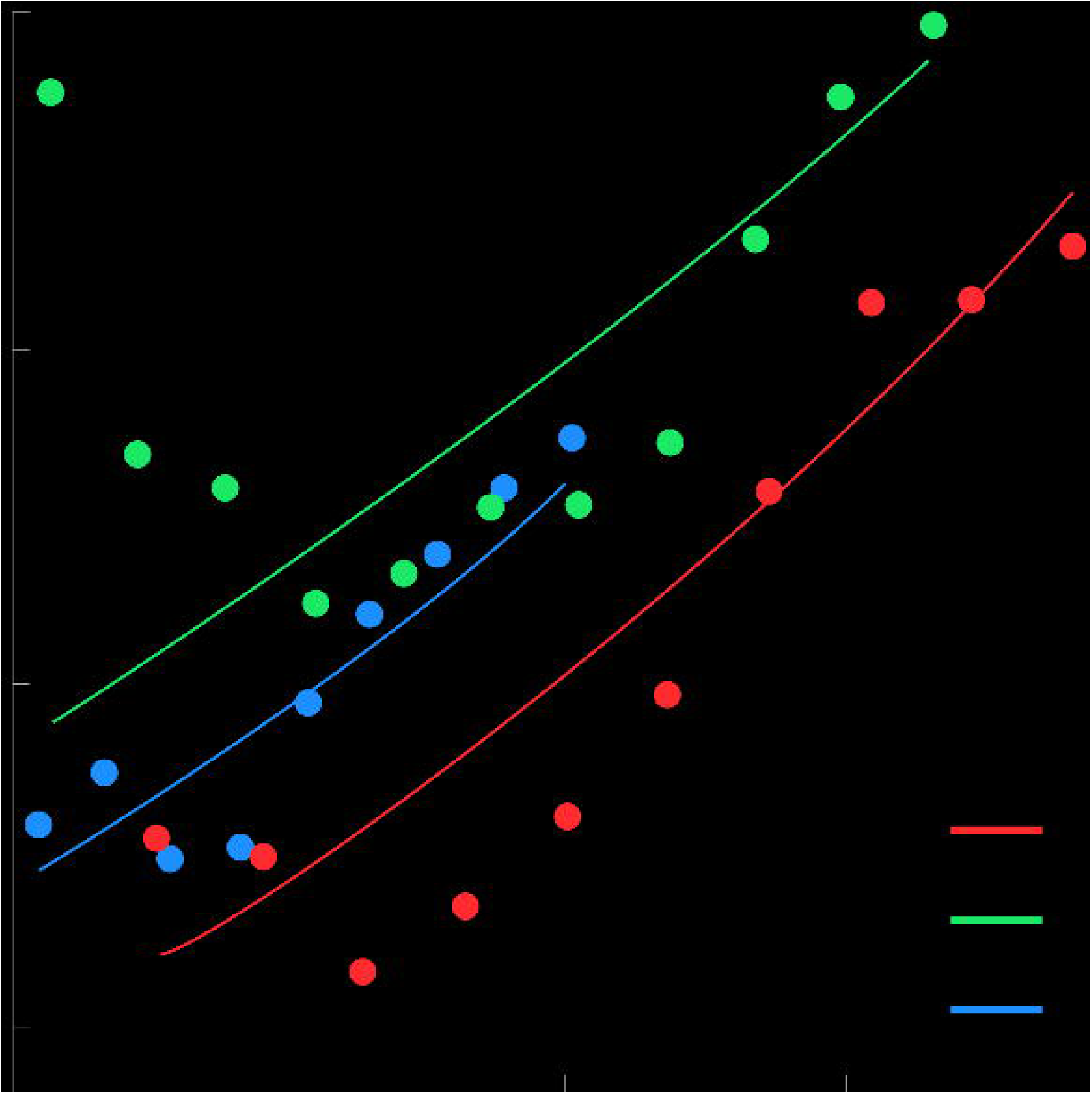
Prey velocity dependent strategy selection. All the subjects consistently show biases using PVBP when the prey velocity is faster. Logistic regression was performed between prey velocity and a categorical dependent variable (0: non-PVBP, 1: PVBP). The p-values of all logistic coefficients were significant (p < 0.001).

